# Fungus causing White-Nose Syndrome in bats accumulates genetic variability in North America and shows no sign of recombination

**DOI:** 10.1101/121038

**Authors:** Jigar Trivedi, Karen Vanderwolf, Vikram Misra, Craig K. R. Willis, John Ratcliffe, James B. Anderson, Linda M. Kohn

**Affiliations:** University of Toronto, Mississauga, ON, Canada.; University of Wisconsin, Madison, WI, USA.; University of Saskatchewan, Saskatoon, SK, Canada.; University of Winnipeg, Winnepeg, MB Canada.

## Abstract

Emerging fungal diseases of wildlife are on the rise worldwide (3) and the best lens on the evolution of the fungal pathogens is population genomics.  Our genome-wide analysis shows that the newly introduced North American population of *Pseudogymnoascus destructans*, the causal agent of White-Nose Syndrome (WNS) in bats, has expanded in size, has begun to accumulate variation through mutation, and presents no evidence as yet for genetic exchange and recombination among individuals.

---

Emerging fungal diseases of wildlife are on the rise worldwide [1] and the best lens on the evolution of the fungal pathogens is population genomics. Our genome-wide analysis shows that the newly introduced North American population of *Pseudogymnoascus destructans*, the causal agent of White-Nose Syndrome (WNS) in bats, has expanded in size, has begun to accumulate variation through mutation, and presents no evidence as yet for genetic exchange and recombination among individuals. DNA Fingerprinting [2] and MultiLocus Sequence Typing (MLST) [3], support the hypothesis that introduction to N. America of *P. destructans* was by a genotype of one mating type from Europe, where the fungus is genotypically diverse and both mating types are found [4]. WNS was first reported in bats at a single location in New York State in 2006 and has since spread to a substantial portion of eastern North America and an outlier location in Washington State [5]. WNS is associated with widespread bat mortality, and has driven one of the most common N. American bat species, *Myotis lucifugus*, and at least one other species, *M. septentrionalis*, to the brink of local extinction in eastern N. America [6]. The causal agent of WNS, the fungus *P. destructans*, is cave-adapted and cold-tolerant, and causes cutaneous infection during hibernation, leading to disruption in bats’ torpor-arousal cycles, and ultimately to the depletion of fat reserves crucial to winter survival [7]. European isolates are infectious and cause cutaneous lesions in captive N. American and European bats, but in Europe mass mortality of bats has not been observed; host and pathogen apparently are able to coexist [7]. This is consistent with the hypothesis that the hardest-hit N. American bat species were naïve hosts with little intrinsic resistance to *P. destructans*.

Our primary question was whether or not the clonal population of *P. destructans* in N. America has begun to accumulate genetic variability through mutation. Given variation, we then asked whether or not the population displays a signature of recombination. The answers to these questions were not accessible through MLST analysis and required whole-genome analysis. We therefore subjected 17 N. American strains of the fungus to whole-genome sequencing and acquired the sequences of 19 others (Supplemental Information). We also sequenced one European strain of the same MLST haplotype that occurs in N. American (8). We then aligned the individual sequence reads to a genome reference sequence (NCBI Accession PRJNA39257). Lastly, we identified all variants in the genome that passed a specific set of filtering criteria, with verification by viewing alignments around each variant site.

Based on five lines of evidence, population-genomic observations fit the expectations for a young and expanding clonal population (Figure 1 and Supplemental Information). 1. Variation is exceedingly rare – only 83 variants (76 SNPs and 7 indels) were discovered among the 36 N. American strains over the 31 MB genome. 2. All of the mutant alleles are infrequent: 75 of 83 variants exist as singletons, one exists in two strains, five exist in three strains, one exists in four strains, and one exists in seven strains. Consistent with this distribution, Tajima’s D, which measures deviations from equilibrium conditions, is strongly negative (-2.7). 3. The base-spectrum of SNPs resembles that of spontaneous mutation, with an excess of C/G to T/A transitions, indicating these mutations are new and not yet influenced by selection (53 of 76 SNPs). 4. Recombination is not detected – none of the pairwise comparisons of the variant (biallelic) sites revealed all four possible genotypes, the criterion of Hudson and Kaplan’s four-gamete test for recombination. 5. While the European strain carried the ancestral allele at each of the 83 variant sites identified among the N. American strains, it was by far the most divergent from the others, with 15,793 variants at other positions scattered throughout the genome. Our European strain is therefore substantially different from the strain that originally founded the N. American population, a difference not evident in the previous MLST data (8).

**Figure 1.**
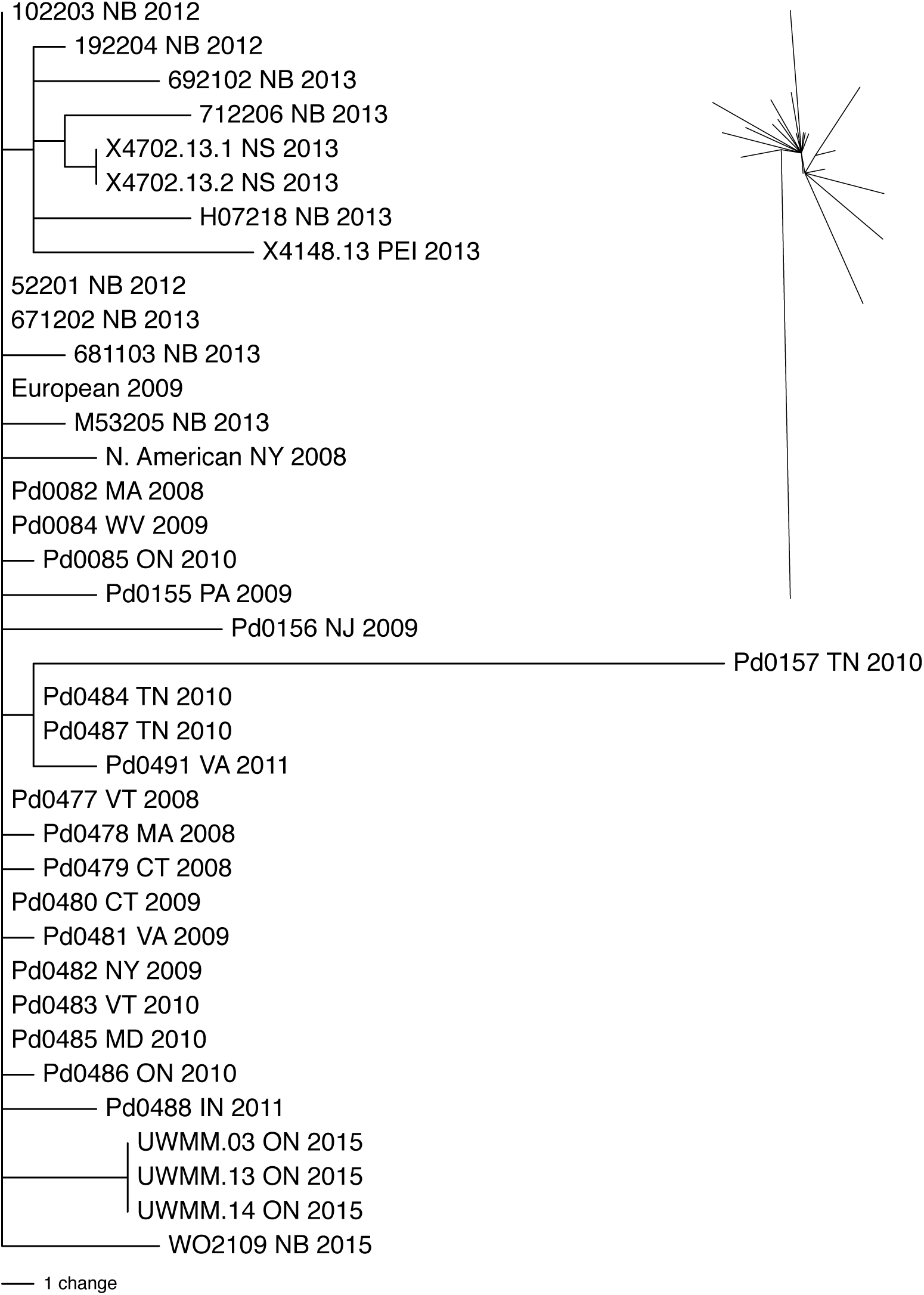
Single most-parsimonious tree of minimum possible length. Left, rectangular tree with strain designations, geographic origin, and date of collection. Top right, unrooted tree without strain labels showing a starburst pattern of diversification.

In parsimony analysis of the 83 variant sites, a single, minimum-length tree (83 steps) of the 37 strains was identified (Figure 1). This tree has no internal conflict (consistency index 1.0); branches therefore represent character-state changes that occurred only once in the tree. The tree illustrates how the mutant alleles, in addition to being rare, are locally distributed. All of the branches leading to strains represent singleton mutations, which by definition occurred in only one place. Also, the strains within the three internal clades defined by non-singleton alleles (represented by internal branches) were geographically restricted within the range of the overall sample.

Our conclusion is that the N. American clone of *P. destructans* has begun to accumulate new alleles through mutation and is in an early stage of diversification. Two questions of critical conservation importance emerge. First, does the fungal population undergo genetic exchange and recombination? Recombination is always a possibility, even in the absence of two mating types and a sexual cycle, because of the well-known capacity of fungi for parasexuality [8]. Given the extreme uniformity of the North American population, individuals of the clone can presumably undergo hyphal anastomosis with mixing of genotypically different nuclei in a common cytoplasm without triggering a somatic incompatibility response. With the accumulation of additional variability in the future, the probability of detecting recombination, if it exists, should go up. Second, how will the further accumulation of variability impact virulence of the fungus on the host bats? The answer here will not only depend on the ongoing population dynamics of the fungus, but also on those of the bats. Evidence suggests that at least one common N. American bat species (*Eptesicus fuscus*) is resistant to, or tolerant of, infection and has apparently not suffered mass mortality [9]. Impacts vary widely for populations of other infected species [6]. Moreover, a slight rebound in *M. lucifugus* populations hit first by WNS, suggests the possibility of some level of increasing resistance in some N. American bat populations [10]. Of particular concern is that new introductions may yet add variability to the N. American population and increase the potential for recombination; such changes in the fungus population could affect the durability and strength of newly appearing resistance in bats.

## Supplemental Information

### P. destructans cultivation and DNA Isolation

*P. destructans* was cultivated on Complete Yeast Medium (CYM, 2 g yeast extract, 2 g peptone, 20 g dextrose, 0.5 g magnesium sulfate, 0.46g of monobasic phosphate buffer, and 1 g of dibasic phosphate buffer per litre) at 10 C. For DNA isolation, strains were allowed to grow for four weeks on sterile cellophane membranes placed on the surface of CYM agar at 10 C. A single colony of each strain was frozen in liquid nitrogen and then lyophilized. Care was taken to ensure that the frozen mycelia did not thaw before completely lyophilization. DNA was isolated by the protocol of Palmer et *al*. (S7). Higher yields of DNA were obtained when the incubation of the mycelia in the lysis buffer was increased from one hour to three hours. The crude DNA preparation was purified with a Qiagen® Gentra Puregene kit.

### Molecular identification of P. destructans

The identity of the fungus was confirmed through ITS gene sequencing. PCR reaction setup consisted of 12.5 μl of 2X mastermix (GoTaq® Green Master Mix, Promega, Madison, WI, USA), 2.5μl each of 5 μM stocks of the ITS1 and ITS4 primers, 6.5 μl of sterile water, and 1μl of the template DNA (original extract diluted 100 fold). PCR reaction conditions were: initial denaturation at 95 C for 2 mins, 35 cycles at 95 C for 30s, 53 C for 30s and 72 C for 1min followed by final extension at 72 C for 10 min. Agarose gel electrophoresis confirmed amplification along with a 100bp molecular marker on a 2% agarose gel in TAE buffer stained with Sybr Safe (ThermoFisher). PCR products were purified by precipitation with PEG before submission for Sanger sequencing at the TCAG sequencing facility.

### Genome sequencing

The genomic DNAs of *P. destructans* were sent to the Centre for Applied Genomics, Sick Childrens Hospital (Toronto, ON) for whole genome sequencing on the Illumina HiSeq platform. After the samples passed the standard quality-control tests (Qubit DNA quantification) for Illumina sequencing, libraries were prepared by the TruSeq Nano protocol (Needs confirmation) and paired-end reads of 151 bp were obtained (Illumina, San Diego, CA, USA). An average of ca. 35 M reads were obtained from each strain for an average coverage depth of 160X. The sequenced strains included the European and North American reference strains (Table S1) as a two-point reference for SNP discovery process for the other North American strains.

**Table S1.** Strains of *P. destructans* used in this study.

| Strain designation | Substrate | Location<sup>b</sup> | Provider<sup>c</sup> | Year | Sequence accession number |
| --- | --- | --- | --- | --- | --- |
| 102203 | *Myotis lucifugus* | White Cave, NB | K. Vanderwolf | 2012 | pending |
| 192204 | *M. lucifugus* | Markhamville Mine, NB | K. Vanderwolf | 2012 | pending |
| 52201 | *M. lucifugus* | White Cave, NB | K. Vanderwolf | 2012 | pending |
| 671202 | *Perimyotis subflavus* | Glebe Mine, NB | K. Vanderwolf | 2013 | pending |
| 681103 | *P. subflavus* | Glebe Mine, NB | K. Vanderwolf | 2013 | pending |
| 692102 | *P. subflavus* | Markhamville Mine, NB | K. Vanderwolf | 2013 | pending |
| 712206 | *P. subflavus* | Markhamville Mine, NB | K. Vanderwolf | 2013 | pending |
| European | *M. myotis* | Thuringa, Germany | V. Misra | 2009 | pending |
| H07218 | *Nelima elegans* | Dorchester Mine, NB | K. Vanderwolf | 2013 | pending |
| M53205 | *Exechiopsis sp.* | Glebe Mine, NB | K. Vanderwolf | 2013 | pending |
| N. American<sup>a</sup> | *M. lucifugus* | NY | V. Mishra | 2008 | pending |
| X4702.13.1 | *M. luciuigus* | NS | K. Vanderwolf | 2013 | pending |
| X4702.13.2 | *M. luciuigus* | NS | K. Vanderwolf | 2013 | pending |
| Pd00082<sup>d</sup> | *M. lucifugus* | MA | J. Foster | 2008 | pending |
| Pd00084 | *M. septentrionalis* | WV | J. Foster | 2009 | pending |
| Pd00085 | *M. lucifugus* | ON | J. Foster | 2010 | pending |
| Pd00155 | *M. lucifugus* | PA | J. Foster | 2009 | pending |
| Pd00156 | *M. lucifugus* | NJ | J. Foster | 2009 | pending |
| Pd00157 | *M. septentrionalis* | TN | J. Foster | 2010 | pending |
| Pd00477 | *M. septentrionalis* | VT | J. Foster | 2008 | pending |
| Pd00478 | *M. septentrionalis* | MA | J. Foster | 2008 | pending |
| Pd00479 | *M. lucifugus* | CT | J. Foster | 2008 | pending |
| Pd00480 | *M. lucifugus* | CT | J. Foster | 2009 | pending |
| Pd00481 | *P. subflavus* | VA | J. Foster | 2009 | pending |
| Pd00482 | *Eptesicus fuscus* | NY | J. Foster | 2009 | pending |
| Pd00483 | *M. lucifugus* | VT | J. Foster | 2010 | pending |
| Pd00484 | *P. subflavus* | TN | J. Foster | 2010 | pending |
| Pd00485 | *M. septentrionalis* | MD | J. Foster | 2010 | pending |
| Pd00486 | *M. lucifugus* | ON | J. Foster | 2010 | pending |
| Pd00487 | *M. septentrionalis* | TN | J. Foster | 2010 | pending |
| Pd00488 | *M. lucifugus* | IN | J. Foster | 2011 | pending |
| Pd00491 | *M. lucifugus* | VA | J. Foster | 2011 | pending |
| UWMM.03 | *M. lucifugus* | Thunder Bay, ON | C. Willis | 2015 | pending |
| UWMM.13 | *M. lucifugus* | Thunder Bay, ON | C. Willis | 2015 | pending |
| UWMM.14 | *M. lucifugus* | Thunder Bay, ON | C. Willis | 2015 | pending |
| WO2109 | Cave Wall | White Cave, NB | K. Vanderwolf | 2015 | pending |
| X4148.13 | *M. lucifugus* | PEI | K. Vanderwolf | 2013 | pending |

<sup>a</sup>Same strain sequenced by the Broad Institute.
<sup>b</sup>Two letter designation for states and provinces (except PEI).
<sup>c</sup>Note: Substrate name was specified by the collector.
<sup>d</sup>For all strains beginning with “Pd” sequence data were generated and supplied by the provider; living representatives were not transferred.

### European origin of N. American P. destructans

A recent MLST study (S2) used eight polymorphic gene loci to identify eight haplotypes in Europe, of which one, ‘Haplotype 1’ was identified as the source of *P. destructans* in North America. The eight sequences of ‘Haplotype 1’ were therefore concatenated and used as a reference for alignment of the Illumina reads of each of the strains in our sample with Geneious 9.1. The genome sequences of all strains in our sample were identical to the Haplotype 1 sequences – no variation was detected in these regions among the 36 North American strains.

### Data pre-processing and read alignment

Raw sequence data (Fastq) of each strain were initially pre-processed with Trimmomatic tool to remove the custom next-gen sequencing adapters/primers from the raw data.

The pre-processed reads were aligned to the *P. destructans* reference genome (*Geomyces destructans* Sequencing Project, Broad Institute of Harvard and MIT, NCBI Accession PRJNA39257) by using Burrows-Wheeler Aligner tool (S3). Overall, around 99.99% of the reads aligned to the *P*. destructans reference genome. A binary version (BAM format) of the SAM file was created by using SortSam command. These files were readied for variant discovery by using Markduplicates and BuildBamIndex functions. These pre-processing functions were part of the Picard tools package <u>(http://picard.sourceforge.net)</u>. To avoid alignment artifacts due to indels and to improve SNP detection, IndelRealigner and RealignerTargetCreator from GATK were used (S1, S4, S9). These functions perform a local realignment of indels resulting in the decrease of false positive SNP calls.

### Variant calling and filtering

We used the GATK tool ‘UnifiedGenotyper’ (v3.5; S4) to identify SNPs and short indels. All 37 strains were run jointly, with ploidy set to two. Given that *P. destructans* is haploid, any site identified as heterozygote indicates potential mapping errors from paralogous loci or DNA sample contamination. When identifying extremely rare variants mapping errors are a common cause of false positives and filtering sites with two haplotypes has proven effective in previous work (S5, S6). We applied the following filters on candidate variants: 1. Genotype quality (‘GQ’) greater than 10; 2. All called individuals were homozygous; 3. Only two alleles present at the site. All filtered candidate variants were visually inspected in the Integrative Genomics Viewer (IGV) (S8) to further eliminate the possibility of erroneous calls. We excluded all variants from the European strain from the analysis.

### Maximum Parsimony analysis

For parsimony analysis, the 37 strains were considered as taxa, the 83 variant sites were considered characters, and their alleles character states. The initial analysis was done by hand; a single most-parsimonious tree of the shortest possible length, 83 steps, and no internal conflict was identified. PAUP4.0 was used to confirm the optimal tree and to generate Figure 1.

